# Amplifying and ameliorating light avoidance in mice with photoreceptor targeting and CGRP sensitization

**DOI:** 10.1101/2025.05.24.655946

**Authors:** Eric A. Kaiser, Audrey Cavanah, Geoffrey K. Aguirre, Frances E. Jensen

**Affiliations:** Department of Neurology, University of Pennsylvania, Philadelphia, PA 19104

**Keywords:** photophobia, migraine, photoreceptors, melanopsin, ipRGCs, CGRP

## Abstract

**Objective:** To determine the photoreceptor basis of light avoidance in mice and assess the effect of CGRP sensitization on this behavior.

**Background:** Prior studies have suggested that photophobia is mediated by a subset of retinal ganglion cells (RGCs) that contain melanopsin, making them intrinsically photosensitive (ipRGCs). These cells also receive extrinsic input from cones, which can also mediate light sensitivity. Here, we examined whether spectral variation targeting melanopsin or specific cone types in mice could effectively model light sensitivity. Also, we assessed whether sensitizing mice with calcitonin gene-related peptide (CGRP) could amplify ipRGC-mediated light avoidance.

**Methods:** Light avoidance behavior was observed in a two-zone chamber illuminated by narrow-band LEDs targeting photopic opsins: 365 nm (UV; rodent S-cone), 460 nm (blue; melanopsin), and 630 nm (red; human L-cone). In a non-targeted assay, we assessed the degree of light avoidance in wildtype C57BL/6J mice to varying intensities (5 to 100%) of the blue and red LEDs. In a targeted assay, mice were given a choice to spend time between zones with differing relative contrast levels (0.50, 0.75, or 1.00) for the targeted photoreceptor(s). This was assessed in two transgenic mice with: 1) human red cone knock-in (RCKI), or 2) adult-onset ablation of M1 ipRGCs (*Opn4^aDTA^).* Mice were studied without intervention or following priming with either peripheral CGRP or vehicle administration every other day for 9 days. A primary measure (mean +/- SEM) was the asymptote value (AV).

**Results:** Wildtype mice showed greater light avoidance with increasing light intensity, demonstrating a parametric response. RCKI mice showed avoidance of the high melanopsin (1.00: 0.52 ± 0.08; n = 18) and L-cone (1.00: 0.30 ± 0.11; n = 15) contrast zones but showed a preference for the higher S-cone (1.00: −0.35 ± 0.06; n = 16) contrast zone. These effects decreased with less relative contrast and, thus, contrast dependent. Adding S-cone contrast opposed avoidance to melanopsin (0.10 ± 0.12; n = 14) or L-cone (−0.19 ± 0.10; n = 15) contrast. Ablation of ipRGCs in *Opn4^aDTA^* mice attenuated avoidance of melanopsin and preference for S-cone stimulation compared to control littermates. On day 9, CGRP priming led to significantly increased avoidance of melanopsin stimulation (0.58 ± 0.08, n = 21) as compared to vehicle priming (0.26 ± 0.09, n = 22) (*F* (1,41) = 5.70, p = 0.02).

**Conclusions:** Our findings further support that ipRGCs play a key role in mediating photophobia. This aversive response to light stems from ipRGCs combining excitatory input from intrinsic melanopsin stimulation and extrinsic L-cone input, which can be opposed by extrinsic inhibitory S-cone input. Chronic exposure to CGRP is likely one of many mechanisms in migraine that can amplify ipRGC signals, leading to photophobia.

**Plain Language Summary:** To better understand light sensitivity, we studied which cells in the eye cause mice to avoid light. We found that mice avoided blue and red light but preferred UV light, and this is the result of a special cell (ipRGCs) in the eye that combines these light signals. Repeated exposure to CGRP, a key nervous system messenger in migraine, increased avoidance of blue light, which may model what happens in people with chronic migraine who experience light sensitivity.

## 1. Introduction

Bright light can be perceived as uncomfortable, and discomfort from light is increased in numerous ophthalmologic and neurologic conditions, most notably migraine headaches. Light sensation begins in the retina, with phototransduction under daylight conditions chiefly by the cones and melanopsin. Mice possess two cone classes (M and S), corresponding to “medium” (green) and “short” (ultraviolet) wavelengths, respectively (Figure 1A). Additionally, a subset of retinal ganglion cells contain the photopigment melanopsin, rendering them intrinsically photosensitive (ipRGCs), with peak sensitivity to cyan wavelengths (Figure 1A).^1, 2^ Like the “classical” RGCs that do not express melanopsin, the ipRGCs (especially the M1 class) also receive extrinsic inputs from the cones.^3^ A topic of general interest is how these different retinal signals contribute to a sense of visual discomfort.

**Figure 1.**
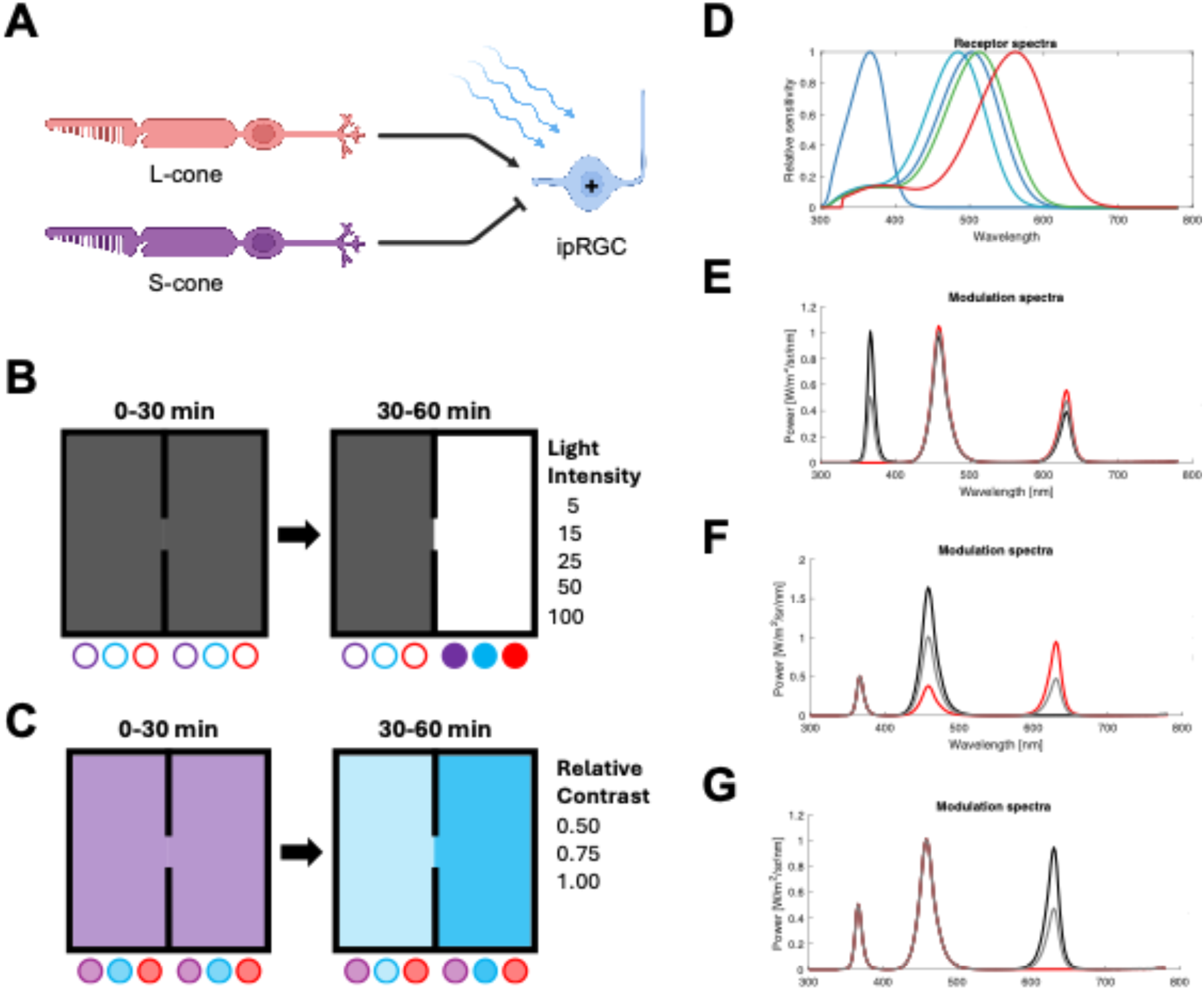
Experimental overview. **A.** Schematic of the retinal network for transgenic red cone knock-in (RCKI) mice, in which L-cone stimulation provides excitatory, extrinsic input to ipRGCs, and S-cones stimulation provides inhibitory, extrinsic input. Melanopsin stimulation provides excitatory, intrinsic input to ipRGCs, which integrate these three signals. **B**. In the non-targeted behavioral assay, animals were placed in the two-zone box. From 0 to 30 min, LEDs were off in both zones. From 30-60 min, LEDs remained off in one zone, while the UV, blue, and UV LEDs were turned on in the second zone. Note that the UV LED was unintentionally blocked by a diffusing panel. The intensity of the LEDs was set to 5, 15, 25, 50, or 100% of their maximal setting. **C.** In the photoreceptor-targeted behavioral assay, animals were placed in the two-zone box. From 0 to 30 min, the LED settings in the two zones were changed to create a difference in contrast upon targeted photoreceptor(s) between the two chambers. The LED settings were scaled so that this stimulus produced 50, 75, or 100% of the maximal contrast available on the targeted photoreceptors. **D**. The spectral sensitivity functions of the relevant photoreceptors under daylight conditions, including mouse S-cone (blue), melanopsin (teal), mouse M-cone (green), human L-cone (red). For panels E-G, shown are three sets of spectral modulations (background: gray, low contrast: red; high contrast: black) used for the photoreceptor-targeted behavior assay. The photoreceptors targeted here include **E)** mouse S-cones; **F**) melanopsin; and **G**). human L-cones.

Prior work in rodents suggests that the ipRGCs are central to the discomfort response. When exposed to blue light, neonatal mice, which have yet to develop functional rods and cones but have functional ipRGCs at birth, exhibit avoidance behaviors^4^ and produce negative vocalizations.^5^ Adult mice show a preference for dark as compared to brightly lit spaces,^6^ and inactivation of the ipRGCs has been shown to attenuate this preference.^7–9^ Notably, the inactivation of the cones also reduces light avoidance,^7^ and recent work indicates that the S-cones and M-cones provide opposing inputs to the ipRGCs that modulate light-aversive behavior.^10^

It appears that these photoreceptor signals mediate aversive behavior, at least in part through the somatosensory trigeminal system. In rats, exposure to very bright light produces neural activation within the trigeminal nucleus caudalis, the subnucleus associated with pain transmission.^11^ Responses to bright light are found in the posterior somatosensory thalamus in neurons that also receive dural afferents.^12^ Optogenetic stimulation of this thalamic nucleus evokes light-aversive behavior in mice.^13^

The interaction of light and trigeminal signals in discomfort is a feature of migraine headaches.^14–16^ Signals from the ipRGCs appear to be amplified in individuals with migraine,^17, 18^ who commonly experience increased light sensitivity during and sometimes even between attacks.^19–27^ An important therapeutic model in migraine has been the induction of light aversive behavior in mice by administration of calcitonin gene-related peptide (CGRP), a key neuropeptide involved in migraine pathophysiology that modulates nociceptive pathways and mediates neurogenic inflammation.^28–31^ An important, unanswered question is if the effect of CGRP is specifically to enhance the aversive behavior evoked by ipRGC signals.

Here, we sought to replicate prior rodent studies by establishing the photoreceptor basis of light-aversive behavior and then extend this paradigm to examine whether CGRP administration amplifies ipRGC signals for light avoidance. We modified a previously described light aversion behavioral assay ^32–35^ to deliver spectral modulations of light that target specific photoreceptors using the technique of silent substitution.^36^ In these studies, the use of a transgenic mouse, in which the mouse M-cone is replaced with the human L-cone, allowed our light modulations to produce larger differential targeting of the photoreceptor signals. We specifically examined aversion to melanopsin and cone stimulation, both in isolation and in combination, and explored if S-cone stimulation could oppose the aversive effects of melanopsin and L-cone stimulation. We used a genetic ablation of melanopsin-containing RGCs to determine if our behavioral observations depended on ipRGC function. Finally, we examined if repeated exposure to CGRP enhanced light avoidance by the ipRGCs. Our findings add photoreceptor specificity to the key translational migraine model of CGRP-induced light avoidance.

## 2. Methods

### 2.1 Animals

Wildtype and transgenic strains of mice used for experiments included: 1) wildtype C57BL/6J mice (Jackson Labs, Bar Harbor, ME); 2) transgenic red cone knock-in mice (RCKI, B6.129-Opn1mw^tm1(OPN1LW)Nat^/J; Jackson Labs, Bar Harbor, ME), and 3) adult-onset genetic ablation of M1 ipRGCs (Opn4^aDTA/aDTA^, *Opn4^tm4.1(DTA)Saha^/J)*. C57BL/6J mice were shipped at 9 weeks of age and then acclimated in our animal facilities for a minimum of 7 days prior to testing. Alternatively, C57BL/6J mice (Jackson Labs, Bar Harbor, ME) were bred, and subsequent litters were also used for testing.

Male and female mice were tested between 10 and 22 weeks of age. All animals were housed in groups of 2 - 5 per cage in standard conditions, on a 12-hour light cycle (on at 0640 ET, off at 1840 ET), and with access to water and food ad libitum. Animal care procedures were approved by the University of Pennsylvania Animal Care and Use Committee and performed in accordance with the standards set by the National Institutes of Health.

### 2.2 Intraperitoneal (ip) drug administration

CGRP was diluted with Dulbecco PBS (Hyclone) as the vehicle. Mice were injected with either 0.1 mg/kg of rat α-CGRP (Sigma) or vehicle, which was administered at 10 μL/g bodyweight with BD ultrafine 31g insulin syringes. Injections were performed by E.A.K. or A.C. just prior to behavioral testing. Animals were gently restrained but not anesthetized during injection.

### 2.3 Light aversion testing chamber

The open field and infrared tracking equipment (Med Associates Inc., St. Albans, VT) have been described.^34^ The field was divided into two equal-sized zones with an opening for free movement between the two zones. Each zone contained a panel of LEDs with three different, narrow-bandwidth LEDs with peak wavelength intensities of: 365 nm (realUV LED Strip Lights, Waveform Lighting), 460 nm (SimpleColor Blue LED Strip Lights, Waveform Lighting), 630 nm (SimpleColor Red LED Strip Lights, Waveform Lighting). Each set of LEDs was digitally controlled. See Supplemental Materials for additional details.

### 2.4 Non-targeted light aversion behavior assay

On the day of the experiment, mice were briefly transported to the testing room and then allowed to acclimate in their home cage to the testing room for at least 1 h with standard overhead fluorescent lighting. All sound-generating equipment was turned on during acclimation and remained on until testing was complete. Behavioral testing was performed between 0900 and 1830 local time. See supplemental material for a description of ambient light conditions.

Following acclimation, mice were placed into one of the zones and allowed to explore both zones freely for 60 min. From 0 to 30 min, both LED panels were off, and then, from 30-60 min, one of the panels was turned on with the blue and red LEDs set to a specific light intensity (between 5% (116 photopic lux, 387.75 photopic lux) and 100% of the maximum (2440 photopic lux, 8624 melanopic lux). The opposing LED panel remained off in a standard light-dark box paradigm. Note that a plexiglass panel had been placed under the LEDs to diffuse light, which was later discovered to block UV light; thus, there was essentially no UV light transmission for this set of experiments.

Animals were tested ∼48 hours later following the same protocol; however, mice were administered CGRP or vehicle as described previously and then immediately placed back in the chamber for 60 min under the same light paradigm described above.

### 2.5 Photoreceptor-targeted light aversion behavior assay

Following acclimation as described in the non-targeted assay, mice were also placed into one of the zones and allowed to freely explore both zones for 60 min. From 0 to 30 min, both LED panels were turned on with the UV, blue, and red LEDs, each set to 50% light intensity. From 30 to 60 min, the LED panels illuminating both zones were adjusted to create a selective difference in contrast upon the targeted photoreceptor(s) using the principles of silent substitution.^36^ The UV, blue, and red LEDs were digitally scaled in each LED panel to produce a stimulus of 50, 75%, or 100% of maximal relative contrast available on the targeted photoreceptor(s) (for examples of spectral modulations used, see Figures 1E-G). The targeted photoreceptor classes included mouse S-cone, melanopsin, and/or human L-cone (see Figure 1D for spectral sensitivity functions). Light flux refers to stimuli in which all three photoreceptors are targeted. The nominal photoreceptor contrasts produced for each spectral modulation at 100% (or 1.00) relative contrast are reported in Table 1. For Opn4^aDTA^ mice, melanopsin-targeted conditions had equal nominal contrast for melanopsin and mouse M-cones (Table 1). Note that the diffusion panel was no longer used for this set of experiments, so UV light was not unintentionally blocked.

**Table 1.** Nominal contrast levels for each photoreceptor type generated by the spectral modulations used for the photoreceptor-targeted light aversion behavior assay.

| **Photoreceptor Direction** | **mouse S-cone** | **mouse Mel** | **mouse Rod** | **mouse M-cone** | **human L-cone** |
| --- | --- | --- | --- | --- | --- |
| **S-cone** | 99.61 | 0.00 | 0.80 | 1.49 | 0.00 |
| **Melanopsin** | 0.00 | 59.88 | 58.97 | 57.55 | 0.00 |
| **L-Cone** | 0.00 | 0.00 | 0.26 | 0.84 | 35.11 |
| **Light Flux** | 100.00 | 100.00 | 100.00 | 100.00 | 100.00 |

### 2.6 CGRP Priming

Modifying a previously described protocol,^37^ animals were administered either ip CGRP or vehicle as described above every other day over 9 days (i.e., days 1, 3, 5, 7, 9). On days 1, 5, and 9, animals were placed in the behavior chamber immediately after administration of CGRP or vehicle and tested as per the photoreceptor-targeted light aversion behavioral assay. On days 3 and 7, animals were returned to their home cages following ip injections.

### 2.5 Statistical Analysis

Data were processed and analyzed using Microsoft Excel (Microsoft Corporation), RStudio (version 2024.090+375, Posit-Software, PBC), and Prism 10 for Mac OS X (Graphpad Software, Inc). Results were reported as mean ± standard error of the mean. Time in the light zone or the high contrast zone for the targeted photoreceptor was reported as 5-min binned intervals over the 60-min testing duration (12 intervals in total). For each interval, time in the light zone (or the high contrast zone) was normalized to the mean time spent in that zone over the first six intervals (0 to 30 min). Asymptote value (AV) was defined as 1 minus the mean normalized time in the light zone (or the high contrast zone) over the final 3 – 5 min intervals (45 to 60 min).

A two-way ANOVA was used to assess the effect of a specific condition (e.g., specific photoreceptor(s) targeted, genotype) over the 60 min testing duration. This test was conducted for the normalized time in light (or high contrast) zone, and subsequent post-hoc analyses were performed to compare differences at a specific 5-min interval using the Sidak multiple comparisons test. A Mann-Whitney test was performed when comparing the mean asymptotic value between two different conditions.

Mice were excluded from the analysis due to equipment or software failure. Mice were excluded for atypical exploratory behavior based on one of two criteria: 1) failing to explore both zones during one of the 5-min intervals during the 0 to 30 min acclimation period or being identified as an outlier using the ROUT method (Q = 1%).

## 3. Results

### 3.1 Light avoidance behavior depends on the contrast level

First, we determined how the degree of light avoidance in a light-dark box changes as the contrast level between the dark zone and the light zone increases. In our non-targeted light avoidance behavioral assay, wildtype (C57BL6/J) mice were observed for 30 min while mice acclimated to the box with LEDs off in both zones. During the testing period, one zone was illuminated by three sets of narrow-bandwidth LEDs with peak wavelength intensities at 365 nm (UV), 460 nm (blue), and 630 nm (red) (Figure 1A); however, in retrospect, we discovered the UV light emission was unintentionally blocked by a plexiglass diffusion panel. The intensity of the light zone was manipulated by increasing the intensity of the LEDs from their maximal setting, 5% to 15%, 25%, 50%, and 100%. Under these testing conditions, mice were observed for an additional 30 min. Light avoidance was assessed by measuring the time in the light zone, which was normalized to the mean time mice spent in that zone during 30 min acclimation period.

With the initiation of 100% light intensity after the 30 min interval, mice (n = 24) spent less and less time in the light zone during the first three intervals (e.g. 35 min (0.98 ± 0.7), 40. min (0.54 ± 0.6), and then during the final three intervals, the time the light zone reached an asymptote (e.g. 50 min (0.36 ± 0.08)) (Figure 2A). We summarized this light avoidance response as an asymptote value (AV), which is 1 minus the mean time in light for intervals 50, 55, and 60 min; thus, values greater than 0 indicated avoidance of the light stimulus and values less than 0 indicate a preference for the light stimulus. For 100% contrast, the AV was 0.67 ± 0.06 (Figure 2B). For 5-50% light intensity, mice also tended to spend less time in the light zone, with each subsequent 5-min interval until reaching a lower asymptote (Figure 2A). However, as the light intensity diminished, mice spent more time in the light zone, thus less avoidance. Examining the AV as a function of light intensity, we observed a parametric response with the most significant degree of change (or slope) occurring between 15% and 50% contrast (Figure 2B). The degree of light avoidance was clearly dependent upon light intensity, with apparent threshold and ceiling effects upon this behavior.

**Figure 2.**
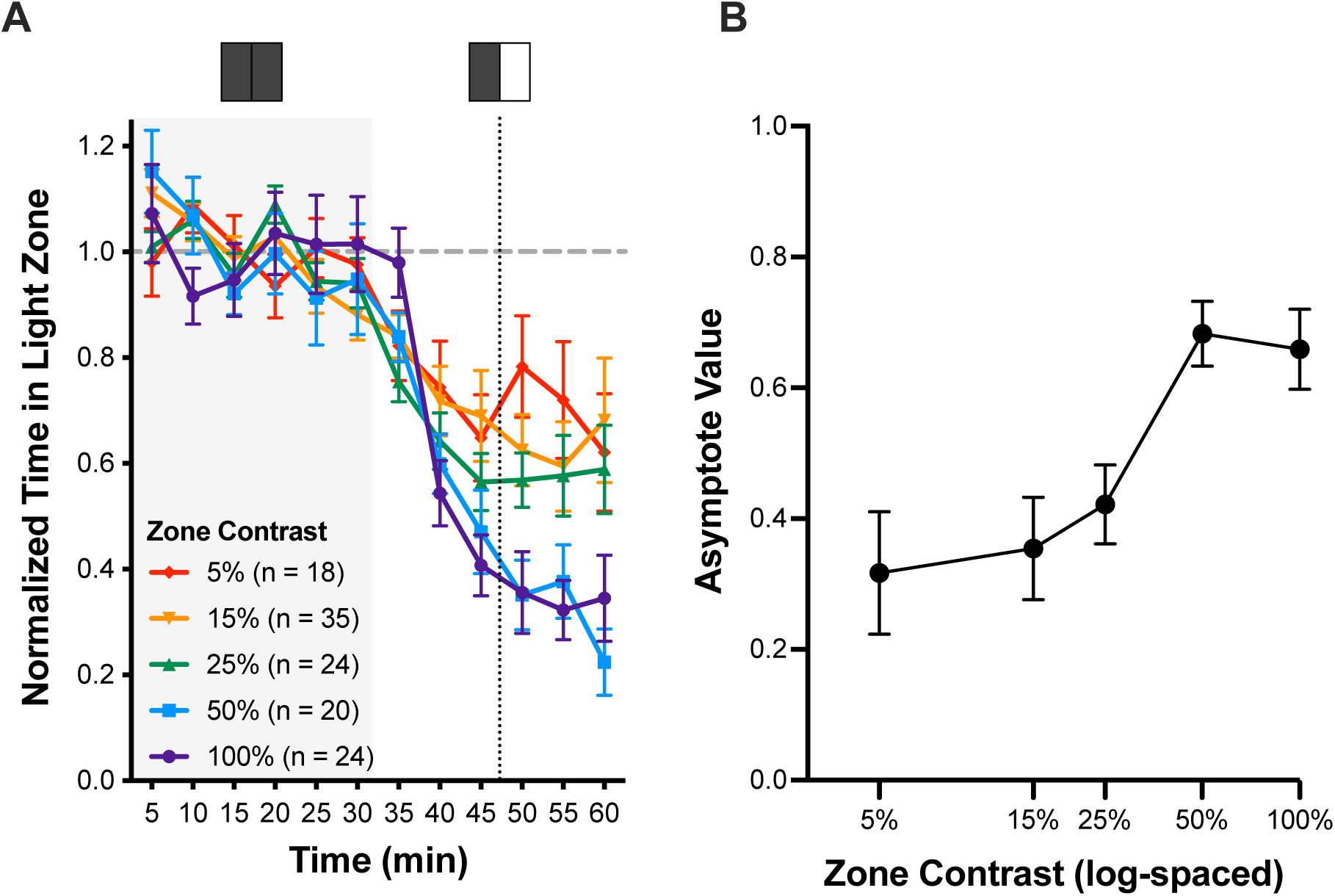
Intensity dependence of light avoidance behavior. **A.** Normalized time spent in the light zone. The light intensities tested were: 1) 5% (red); 2) 15% (orange); 3) 25% (green); 4) 50% (blue); 5) 100% (purple). Wildtype C57BL6/J mice were tested under non-targeted conditions. **B.** Using the asymptote value, light avoidance behavior is expressed as a function of log-spaced zone intensity. The asymptote value is calculated from 1 minus the mean normalized time in light during the 50-, 55-, and 60-min intervals. Error bars indicate mean ± SEM.

### 3.2 Melanopsin and L-cone stimulation elicit light avoidance

To target specific photoreceptor classes, we modified the light avoidance behavioral assay (Figure 1B). From 0-30 min, mice were acclimated to the box with equivocal background light in both zones, for which all three sets of LEDs were set to 50% of their maximal setting.

During the testing period (30-60 min), the UV, blue, and red LEDs were individually adjusted to target specific photoreceptor combinations using silent substitution.^36^ Note that the diffusion panel had been removed, so UV LEDs were no longer unintentionally blocked. One zone was set to a high contrast targeting the specified photoreceptor, and the other zone was set to low contrast. Therefore, in this operant assay, mice could choose whether to spend time in the high or low contrast zone. To better isolate melanopsin versus cone function, we tested transgenic mice with a red cone knock-in (RCKI, B6.129-Opn1mw^tm1(OPN1LW)Nat^/J), replacing the mouse M-cone and thus shifting the spectral sensitivity of that cone into the red range of the visual spectrum.

To establish the light avoidance behavior under these modified conditions, we examined whether mice avoided 100% contrast of UV, blue, and red LEDs, stimulating mouse S-cone, human L-cone, and melanopsin, which was referred to as light flux. Mirroring our observation to 100% contrast in our non-targeted conditions (Figure 2), RCKI showed robust light avoidance to light flux (AV: 0.72 ± 0.05) (Figure 3A, E). Targeted stimulation of melanopsin at 100% also led to light avoidance (AV: 0.52 ± 0.08) (Figure 3B, E). The degree of light avoidance of melanopsin decreased with lower contrast (AV: 0.75: 0.26 ± 0.11; 0.50: 0.05 ± 0.7), demonstrating a contrast-dependent effect. Targeted stimulation of L-cone at 100% contrast also led to light avoidance (AV: 0.30 ± 0.11), although this effect was less robust than melanopsin (Figure 3C, E). Like melanopsin, this response was contrast-dependent. Avoidance to L-cone stimulation also decreased with 0.75 contrast (AV: 0.25 ± 0.11), and avoidance was no longer observed at 0.50 contrast (AV: −0.08 ± 0.08). Overall, melanopsin and L-cone stimulation in isolation and in combination elicited light avoidance behavior.

**Figure 3.**
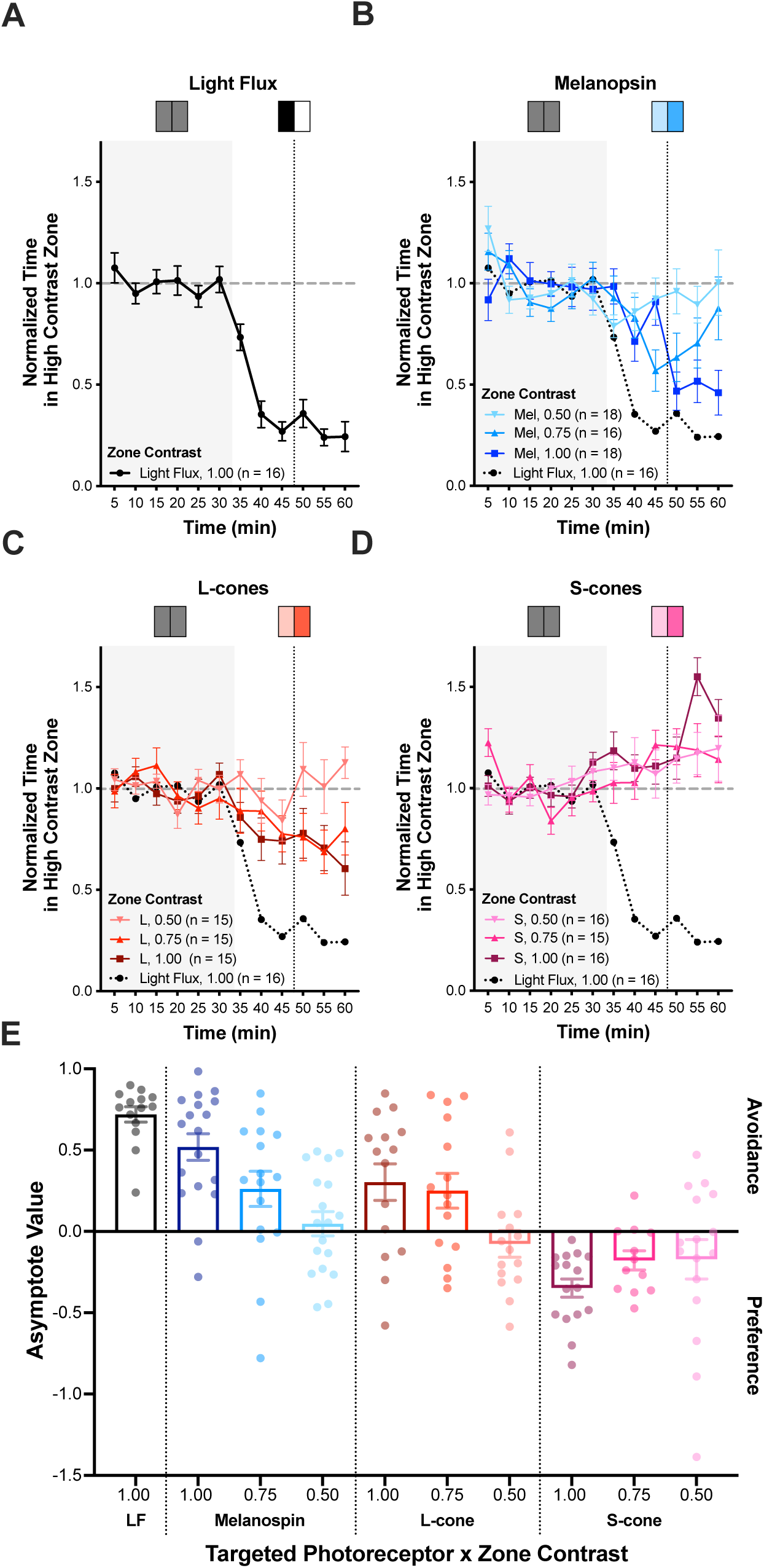
Effect of targeted photoreceptor stimulation on light avoidance or preference behavior. For panels A-D: Normalized time spent in the high contrast zone. The targeted photoreceptors included: **A**. light flux, **B.** melanopsin, **C.** human L-cone, and **D.** mouse S-cone. RCKI mice were tested under targeted photoreceptor conditions. Relative contrast levels between zones were 0.50, 0.75, and 1.00. The effect of light flux (dotted line) is included as a reference in panels B-D. **E.** Asymptote value (AV) for each photoreceptor targeted across each zone contrast. Targeted photoreceptors tested include: 1) light flux (black), 2) melanopsin: 0.50 (light blue), 0.75 (medium blue), 1.00 (dark blue), 3) human L-cone: 0.50 (light red), 0.75 (medium red), 1.00 (dark red), 4) mouse S-cone: 0.50 (light purple), 0.75 (medium purple), 1.00 (dark purple). AV greater than 0 indicates avoidance of the targeted photoreceptor, whereas AV less than 0 indicates preference behavior for the targeted photoreceptor. Plot symbols correspond to individual animals; bars indicate mean ± SEM.

### 3.3 S-cone stimulation elicits light preference

Based on prior observation that S-cones oppose avoidance to L-cones^10^ and provide inhibitory extrinsic input on ipRGCs,^38^ we examined the behavior responses to targeted S-cone stimulation. Here, we observed that RCKI spent more time in the high-contrast zone with S-cone stimulation, which tended to increase over the testing period (Figure 3D). This indicated that mice prefer S-cone stimulation. This light preference behavior was contrast-dependent such that lower levels of S-cone contrast led to less light preference behavior (AV: 1.00: −0.35 ± 0.06; 0.75: y0.18 ± 0.06; 0.50: −0.17 ± 0.12) (Figure 3E). Thus, S-cone stimulation had the opposite behavior response to melanopsin and L-cone stimulation.

### 3.3 S-cone stimulation opposes melanopsin and L-cone light avoidance

To examine this inhibitory effect further, we examined whether adding S-cone stimulation could reduce the light avoidance to melanopsin or L-cone stimulation. Combining S-cone stimulation with melanopsin stimulation led to minimal light avoidance in RCKI mice (AV: 0.10 ± 0.12), which was significantly less than melanopsin stimulation alone (p = 0.009) (Figures 4, S1A). The resulting effect appeared to be nearly additive (e.g. 0.52 (Mel, 1.00) – 0.35 (S, 1.00) = 0.17). Furthermore, adding S-cone stimulation to L-cone stimulation overcame avoidance of L-cone stimulation and led to light preference (AV: −0.19 ± 0.10). This change significantly differed from L-cone stimulation alone (p = 0.004) (Figure 4, S1B). The resulting effect also appeared to be approximately additive (e.g. 0.25 (L, 1.00) – 0.35 (S, 1.00) = −0.10).

**Figure 4.**
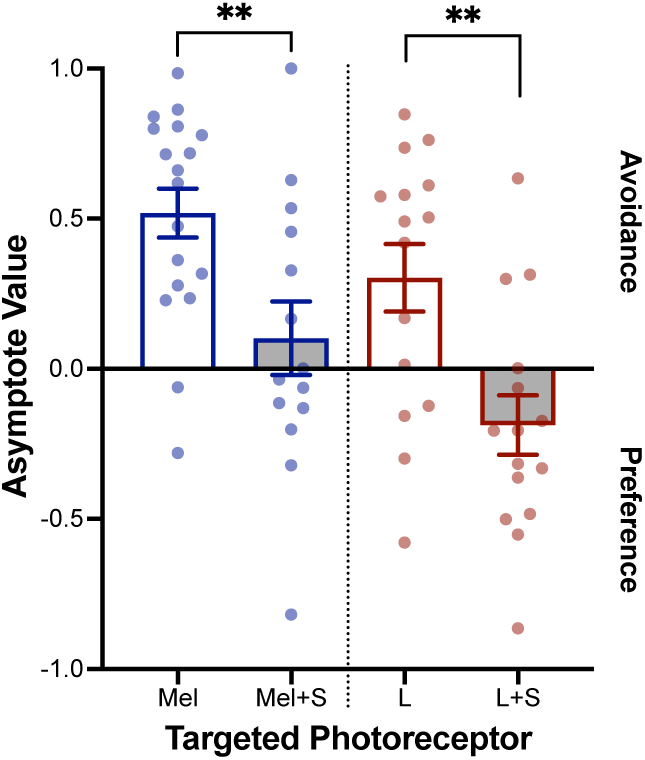
Effect of combing S-cone stimulation to melanopsin or human L-cone stimulation on light avoidance or preference behavior. Asymptote value (AV) for each photoreceptor targeted across each zone contrast. Targeted photoreceptors tested include: 1) melanopsin alone (blue, empty bar), n = 18; 2) melanopsin and mouse S-cone (blue, gray bar), n = 14; 3) human L-cone alone (red, empty bar), n = 15; 4) human L-cone and mouse S-cone (red, gray bar), n = 15. AV greater than 0 indicates avoidance of the targeted photoreceptor, whereas AV less than 0 indicates preference behavior for the targeted photoreceptor. Plot symbols correspond to individual animals; bars indicate mean ± SEM. **p < 0.01.

These findings are consistent with an S-cone inhibitory input on ipRGCs that opposes the excitatory, intrinsic signals from melanopsin and the excitatory, extrinsic input from L-cones.

### 3.4 ipRGCs are key mediators of light avoidance/preference behavior

To assess whether ipRGCs are responsible for mediating light avoidance/preference behavior, we observed these behaviors in transgenic mice that have adult-onset genetic ablation of melanopsin-expressing ipRGCs (Opn4^aDTA/aDTA^ or Opn4^aDTA^) and compared their responses to their control littermates (Opn4^+/+^ or Opn4^WT^). In response to melanopsin-directed stimulation, Opn4^aDTA^ mice (AV: 0.18 ± 0.7) showed significantly less light avoidance as compared to Opn4^WT^ mice (AV: 0.44 ± 0.7; p = 0.007) (Figure 5A, C). When examining the behavior over the 60-min period, genotype had a significant effect (F (1,25) = 15.95, p < 0.001). In response to S-cone stimulation, Opn4^aDTA^ mice (AV: −0.23 ± 0.6) showed less light preference as compared to Opn4^WT^ mice (AV: −0.59 ± 0.18) (Figure 5B, C); however, this reduced effect did not reach statical significance (p = 0.10). Similarly, when examining the behavior over the 60-min period, the genotype did not quite reach significance (F (1,26) = 4.06, p = 0.054). While the likely partial genetic ablation of ipRGCs did not entirely block these responses, these findings suggest that ipRGCs likely play a pivotal role in transmitting melanopsin and S-cone signals to the subcortical and cortical circuits that elicit light avoidance/preference behavior.

**Figure 5.**
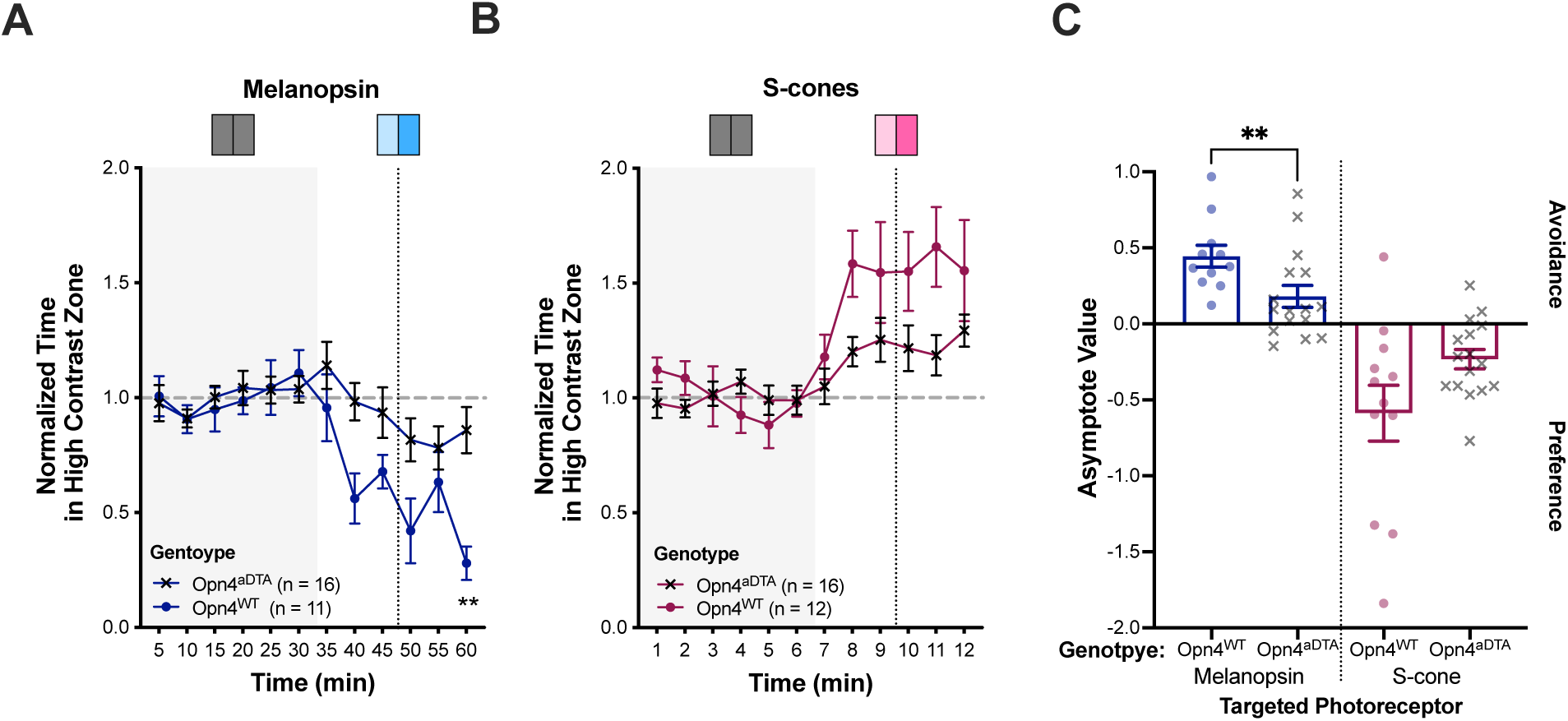
Effect of ablating melanopsin-containing RGCs (ipRGCs) on light avoidance to melanopsin stimulation or preference to mouse S-cone stimulation. For panels A-B: Normalized time spent in the high contrast zone. The targeted photoreceptors included: **A**. melanopsin (dark blue); **B.** mouse S-cone (dark purple). Opn4^aDTA^ mice (n = 16, x symbol) with adult-onset ablation ipRGCs and their control littermates, Opn4^WT^ (n(melanopsin)= 11; n(mouse S-cone) = 12, closed circle) were tested under targeted photoreceptor conditions. The relative contrast level between zones was 1.00. **C.** Asymptote value (AV) for each photoreceptor and genotype as outlined in panel A-B. Each dot represents an individual animal. AV greater than 0 indicates avoidance of the targeted photoreceptor, whereas AV less than 0 indicates preference behavior for the targeted photoreceptor. Plot symbols correspond to individual animals; bars indicate mean ± SEM. **p < 0.01.

### 3.5 CGRP enhances melanopsin-mediated light avoidance

Mason et al. found that wildtype (WT) mice show greater light avoidance following a single administration of ip CGRP as compared to vehicle (Veh);^35^ however, it required very bright white light to induce the behavior. Here, we examined whether CGRP could increase light avoidance in the non-targeted light avoidance behavior assay. We tested C57BL6/J mice two days after their initial testing, as described in Figure 2. We administered either ip CGRP (0.1 mg/kg) or ip vehicle (Veh) to WT mice just before the acclimation period (Figure S2A-B). CGRP-treated animals tended to have greater light avoidance at 15, 50, and 100% contrast than vehicle animals; however, these effects were non-significant.

Next, we examined whether repeated administration of CGRP, potentially inducing priming, could increase the light avoidance of melanopsin stimulation. Here, RCKI mice were administered either ip CGRP (0.1 mg/kg) or Veh every other day over a nine-day period.

Following treatment on days 1, 5, and 9, light avoidance of melanopsin at 0.75 contrast was examined. On day 1, light avoidance of melanopsin was similar between CGRP (AV: 0.57 ± 0.8) and Veh-treated (AV: 0.50 ± 0.07) mice (Figure 6A, C). In Veh-treated mice, light avoidance decreased on day 5 (AV: 0.30 ± 0.09) (Figure S3) and remained stable on day 9 (AV: 0.26 ± 0.09). This mirrors prior observations that mice acclimate to the light avoidance behavioral assay with repeated testing.^34, 35^ In CGRP-treated mice, light avoidance also decreased on day 5 (AV: 0.28 ± 0.11) but increased on day 9 (AV: 0.58 ± 0.08). When examining the behavior over the 60 min testing period on day 9, treatment had a significant effect (*F* (1,41) = 5.70, p = 0.02) with CGRP-primed animals spending less time in the high melanopsin contrast zone than Veh-treated animals starting at the 40 min interval. Per analysis of the AV on day 9, CGRP- primed mice showed significantly greater light avoidance of melanopsin than Veh-primed mice (p = 0.02) (Figure 6C). This suggests that priming with CGRP can enhance light avoidance of melanopsin stimulation.

**Figure 6.**
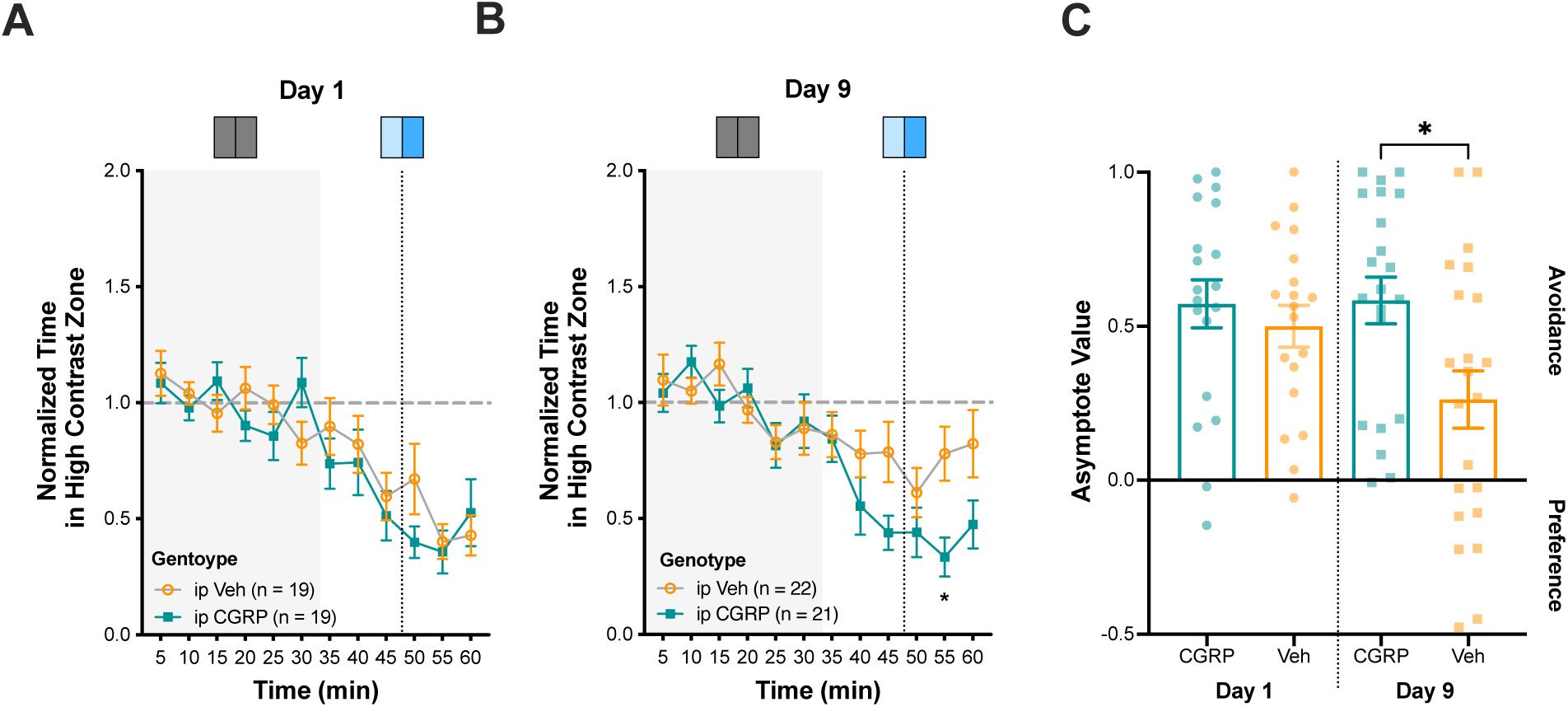
Effect of repeated administration of CGRP on light avoidance behavior to melanopsin stimulation. For panels A-B: Normalized time spent in the high contrast zone with the melanopsin as the targeted photoreceptor at a relative contrast level of 0.75. RCKI mice were administered ip CGRP (0.1 mg/kg) or vehicle (Veh) on days 1, 3, 5, 7, and 9, with testing occurring on day 1 (**A**), day 5 (see Figure S3), and day 9 (**B**). Errors bars indicate ±SEM. **C.** Asymptote value (AV) for each photoreceptor and genotype as outlined in panel A-B. The groups tested included ip Veh and ip CGRP. Each dot represents an individual animal. AV greater than 0 indicates avoidance of the targeted photoreceptor, whereas AV less than 0 indicates preference behavior for the targeted photoreceptor. Plot symbols correspond to individual animals; bars indicate mean ± SEM. *p < 0.05.

## 4. Discussion

Our study examined how light avoidance varies by photoreceptor target and stimulus contrast, how these behaviors are attenuated by ablation of the ipRGCs, and how they are amplified by a CGRP priming procedure. We observed light avoidance behavior in the mouse is sensitive to relatively small differences in light contrast, following a lawful stimulus-response relationship. Similar to prior human work,^10^ we find that the pattern of response to different photoreceptor combinations implicates the ipRGCs as the source of the light avoidance signal. Further support for this proposition was found in the effect of ablating ipRGCs upon light avoidance. A significant additional contribution of this study was our demonstration that CGRP priming amplifies the effect of these photoreceptor manipulations upon light avoidance behavior. The results provided novel evidence of photoreceptor-specific light avoidance responses in a pre-clinical migraine model.

As has been seen previously,^4, 5, 10^ we found that mice avoid the testing chamber that contains a light spectrum enriched for melanopsin and L-cone stimulation. In contrast, increasing S-cone stimulation induced a zone preference. The opposite effects of L- and S-cone stimulation upon behavior are consistent with the opponent synaptic inputs derived from these cone classes upon the M1 class of ipRGCs.^38^ We also found that adult-onset genetic ablation of ipRGCs produced a partial reduction in avoidance of stimulation of melanopsin and long wavelength cones. This ablation procedure is thought to principally inactivate the M1 ipRGC class.^3, 39^ Thus, we feel it is likely the M1 class are a necessary and sufficient signal for maximal light avoidance, which can be downregulated by S-cone input.

Our study made use of silent substitution approaches to target photoreceptor classes. As discussed below, this approach has some limits in the strength of photoreceptor stimulus that may be produced. A distinct advantage, however, is that silent substitution stimuli may also be used to study the intact human visual system, thereby increasing the translational relevance of this rodent work. Indeed, our results may be related to prior studies in humans that have shown that selective melanopsin and cone stimulation produce visual discomfort in a manner suggestive of the integration of these signals in the ipRGCs.^17^ This prior study did not examine isolated S-cone stimulation; thus, it remains unknown if this manipulation may be used to decrease visual discomfort. Support for this idea, however, is found in studies of the human pupil response (which is also mediated by M1 ipRGC signals)^3^ in which isolated S-cone stimulation produces a paradoxical, opponent dilation of the pupil.^40^

An important contribution of our work is demonstrating the photoreceptor basis of light avoidance following CGRP administration. Both peripheral and central CGRP administration can induce light aversive behavior, likely by independent mechanisms.^32–35^ In wildtype mice, we found that CGRP treatment produced greater light avoidance than vehicle-treated mice, but only at high levels of contrast. The vehicle effect here was greater than observed by Mason et al.; this may reflect that mice in this study were not exposed to the chamber twice prior to receiving CGRP or vehicle, thus reducing the exploratory drive of Veh-treated animals. This is the first observation that CGRP priming leads to increased light aversion; this was in response to melanopsin stimulation, directly implicating a critical role for ipRGCs in CGRP-mediated modulation of light aversion.

Given our results, future studies could examine the specific mechanisms whereby CGRP administration enhances ipRGC signals. A plausible mechanism is that peripheral CGRP activates peripheral trigeminal afferents. CGRP priming led to periorbital allodynia in mice by day 5 of the protocol.^37^ Here, we did not observe an effect until day 9, suggesting that modulation of ipRGC signals requires greater central sensitization, given the presence of second-order trigeminal neurons in the trigeminal nucleus caudalis and other subcortical circuits. Similar to the findings in the current study, trigeminal afferent activation by corneal inflammation has led to light aversive behavior in mice, which was dependent on ipRGCs.^41^ Nevertheless, additional studies will need to dissect the neural circuits in which CGRP-induced activation of trigeminal afferents modulate aversive ipRGC signals.

The temporal course of sensitizing ipRGC signals warrants further investigation. We observed the effect of CGRP priming develop over a 9-day course. It is entirely possible that this dynamic process had not reached the asymptotic maximum and that further exposure to CGRP could produce still greater amplification of aversive ipRGC signals. CGRP priming may lead to a number of neuroplastic changes in subcortical pathways, including upregulation of glutamatergic signaling, satellite glia activation, and synaptic modifications of interneurons. The short and long-term mechanistic changes that accompany the development of increased light avoidance may unveil potential pharmacologic targets.

The magnitude of the light avoidance effects we observed was smaller than has been reported in other studies.^32–35^ Our use of silent substitution stimuli is a likely explanation. Non-selective increases in broadband light from darkness can produce massive increases in photoreceptor contrast. Such a stimulus drives all intact photoreceptor classes simultaneously and is arguably more reflective of ecological contexts but does not allow for inferences regarding differential photoreceptor contributions. Here, we accepted the stimulus intensity limitations of the silent substitution approach, given its desirable inferential and translational properties.

## 5. Conclusions

ipRGCs play a key role in the perception of light as aversive in humans and mice, integrating the excitatory input from intrinsic melanopsin stimulation and extrinsic long-wavelength cone input with inhibitory extrinsic input from S cones. Mice appear to be sufficient models of light sensitivity, allowing for future dissection of the downstream neural circuitry. CGRP-mediated central sensitization of the trigeminal system is likely one of multiple neural processes that can modulate ipRGC signals, leading to increased light sensitivity.

---

**Figure S1.**
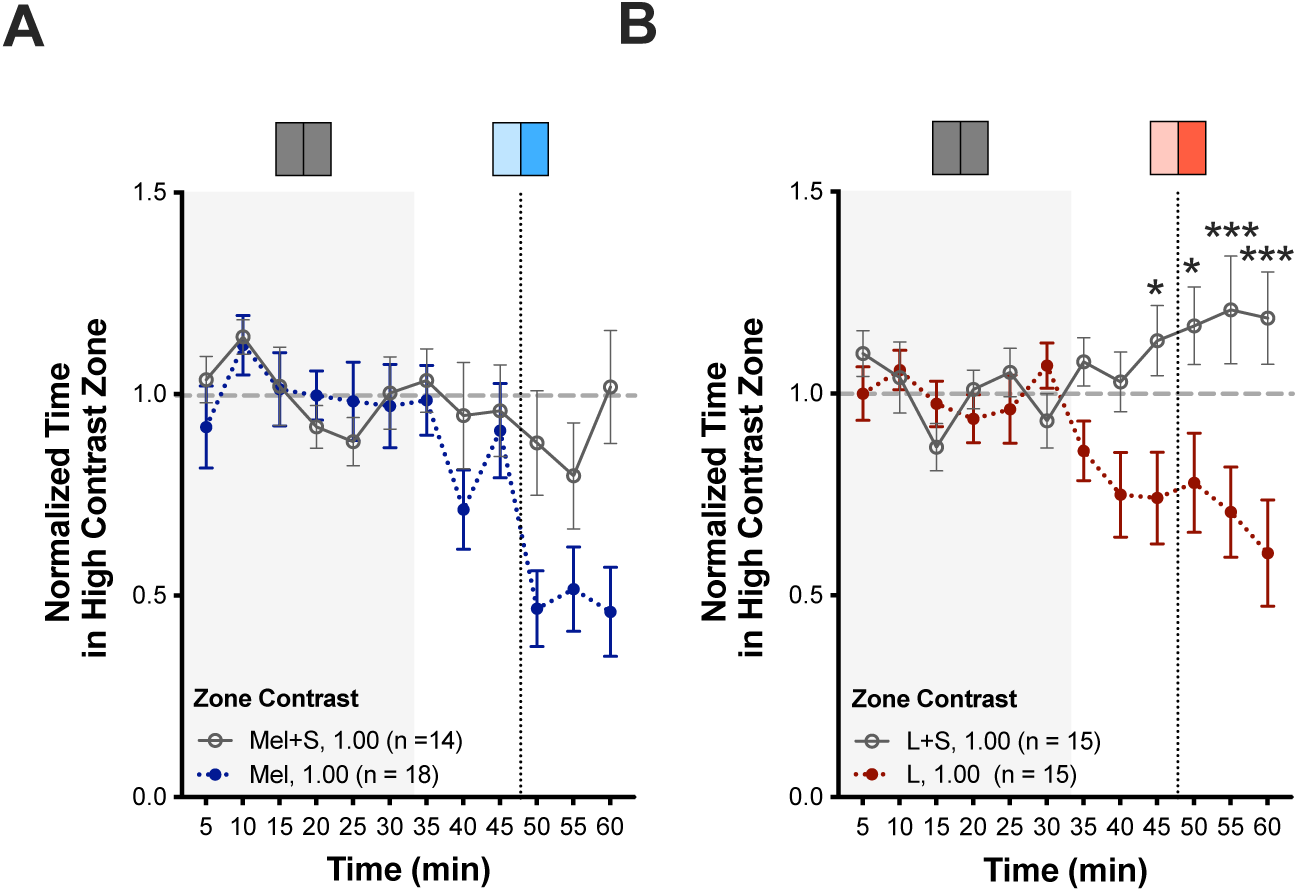
Effect of combing S-cone stimulation to melanopsin or human L-cone stimulation on light avoidance or preference behavior. For panels A-B: Normalized time spent in the high contrast zone. The targeted photoreceptors included: **A**. melanopsin +/- S-cone **B.** human L-cone +/- S-cone. RCKI mice were tested under targeted photoreceptor conditions including: 1) melanopsin alone (dotted dark blue line, closed circle), n = 18; 2) melanopsin and mouse S-cone (gray line, open circle), n = 14; 3) human L-cone alone (dotted red line, closed circle), n = 15; 4) human L-cone and mouse S-cone (gray line, open circle), n = 15. The relative contrast level between zones was 1.00. Error bars indicate mean ± SEM. *p < 0.05. ***p < 0.001.

**Figure S2.**
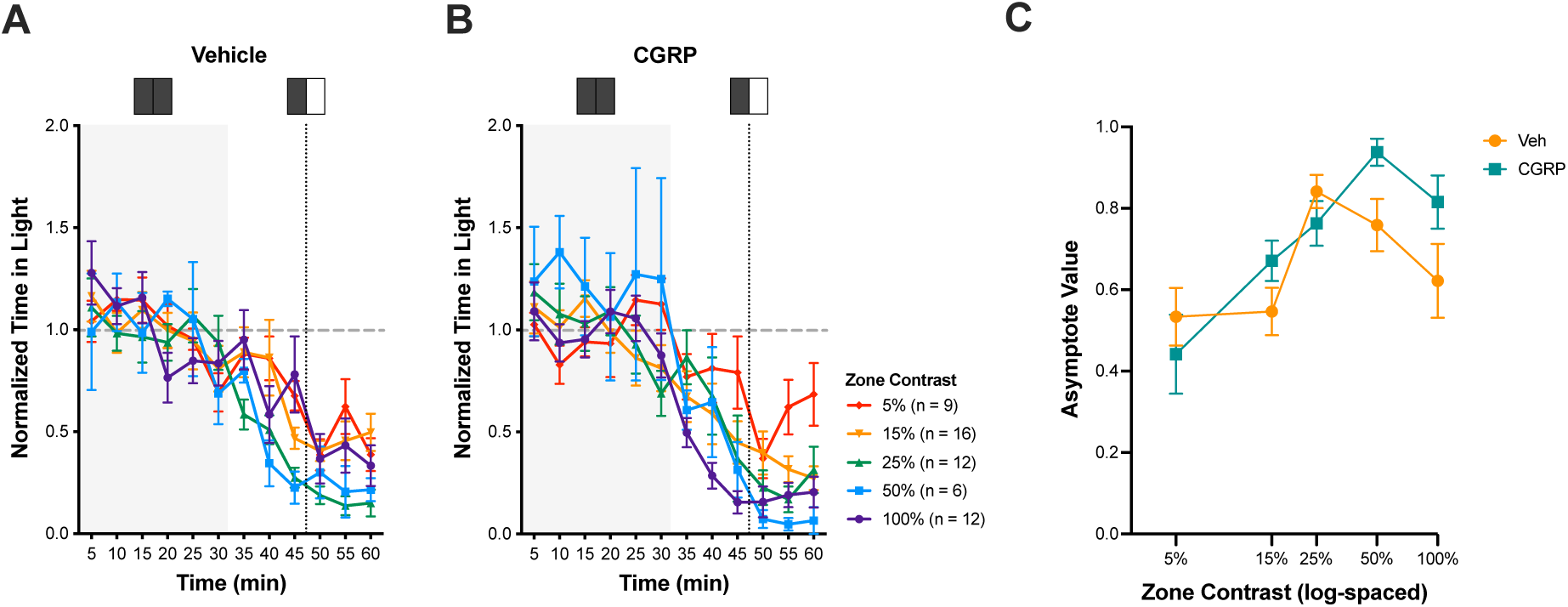
Effect of a single administration of CGRP on light avoidance behavior to non-targeted light stimulation. In panel A-B, normalized time spent in the light zone. The zone contrasts tested were: 1) 5% (red), n = 9; 2) 15% (orange), n(Veh) = 14, n(CGRP) = 16; 3) 25% (green), n = 12; 4) 50% (blue), n = 6; 5) 100% (purple), n = 12. Wildtype C57BL6/J mice were tested under non-targeted conditions. Prior to testing, animals were administered either ip Vehicle (**A**) or CGRP (0.1 mg/kg) (**B**). **C.** Using the asymptote value, light avoidance behavior is expressed as a function of log-spaced zone contrast across groups: 1) ip Vehicle (Veh, orange) or ip CGRP (teal). Error bars indicate mean ± SEM.

**Figure S3.**
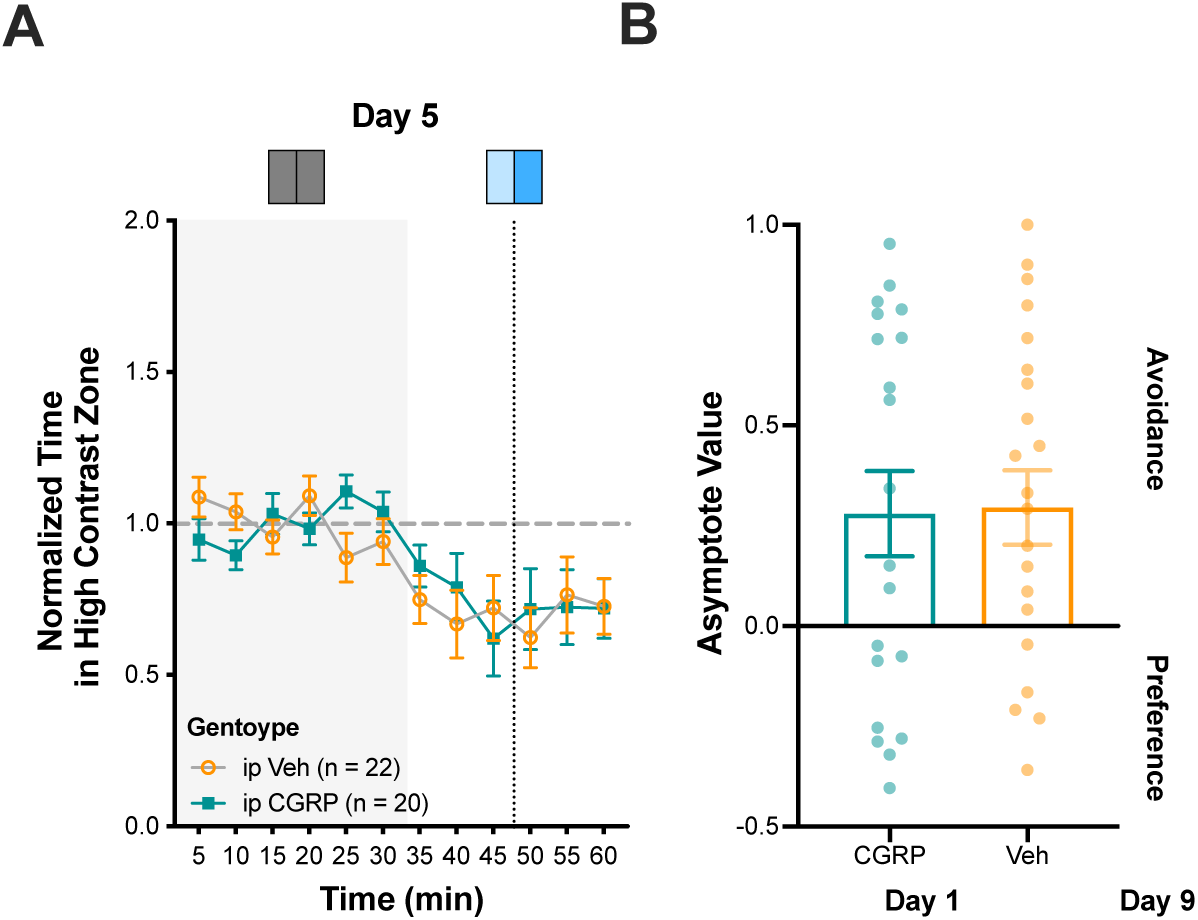
Effect of repeated administration of CGRP on light avoidance behavior to melanopsin stimulation on day 5. **A.** Normalized time spent in the high contrast zone with the melanopsin as the targeted photoreceptor at a relative contrast level of 0.75. RCKI mice were administered ip CGRP (0.1 mg/kg) or vehicle (Veh) on days 1, 3, 5, 7, and 9, with testing occurring on day 1 (see Figure 6A), day 5 (A), and day 9 (see Figured 6B). **C.** Asymptote value (AV) for each photoreceptor and genotype as outlined in panel A. For day 5, the groups tested included ip Veh (n = 22) and ip CGRP (n = 20). Each dot represents an individual animal. AV greater than 0 indicates avoidance of the targeted photoreceptor, whereas AV less than 0 indicates preference behavior for the targeted photoreceptor. Error bars indicate mean ± SEM.

## Supplemental Materials

### S1. Methods

#### S1.1 Ambient Lighting Conditions

With lights on in the housing room, the ambient lighting was 780 photopic lux and 320 melanopic lux, but this ranged from 0.812 to 516 photopic lux and 0.042 to 182 melanopic lux within a cage, depending on the position on the rack. For the room where behavior was assessed, ambient light levels were 451 photopic lux and 222 melanopic lux, but light levels decreased to 57 to 155 photopic lux and 22 to 66 melanopic lux within cages during acclimation.

#### S1.2 Light aversion testing chamber – additional details

Behavior was assessed in a plexiglass open field (27.0 cm wide x 27.0 cm deep x 20.3 cm high) containing two sets of 16-beam infrared arrays (Med Associates Inc., St. Albans, VT). The field was divided into two equal-sized zones by a custom-made acrylic insert that was opaque to visual light but transmitted infrared light (ePlastics, San Diego, CA). The insert was a five-sided, opaque, dark-colored box with a top but no floor. In the center, a removable divider split the chamber into two equal zones and contained an opening (5.2 cm x 6.8 cm), allowing free movement between zones. Each zone contained two vents, one containing a 25 mm x 10 mm, 5 V fan to allow unrestricted airflow and reduce heat generated by the LED panels. A baffle covered the vents to reduce light penetration into the box. Finally, the roof of the box contained a panel of LEDs over each zone. Each testing chamber was enclosed in a sound-attenuating cubicle (56 cm wide x 38 cm deep x 36 cm high), which contained a fan for ventilation (Med Associates Inc.).

Each LED panel was connected to a control box that provided power to the lights, as well as a 32-bit ARM core microcontroller (Arduino Due, Arduino). This allowed for independent, 12-bit digital control of the intensity of the UV, blue, and red LEDs in each panel. Custom MATLAB (The MathWorks, Inc, Natick Massachusetts) code was used to set and change light intensity.

We performed a calibration of the LED panel using a photometer (SpectraScan Spectroadiometer PR-670, JADAK, North Syracuse, NY), measuring the spectral emissions of the blue and red LEDs and confirming a linear gamma function. This photometer could not directly measure UV light spectral emissions, so we relied on tabular values provided by the LED manufacturer to extrapolate a linear gamma function based on our findings from the red and blue LEDs.

Infrared tracking of mouse movement was recorded and analyzed by using Activity Monitor v6.02 (Med Associated Inc.) software.

We measured a 2° C difference in temperature between the chambers under dark and 100% light intensity conditions. Prior work has indicated that this temperature difference is unlikely to change rodent chamber preference.^42^

## Abbreviations

ANOVA: analysis of variance
AV: asymptote value
CGRP: calcitonin gene- related peptide
ipRGCs: intrinsically photosensitive retinal ganglion cells
L: long wavelength
LEDs: light emitting diodes
M: medium wavelength
Mel: melanopsin
RCKI: red cone knock-in
RGCs: retinal ganglion cells
S: short wavelength
UV: ultraviolet
Veh: vehicle
WT: wildtype.

## Acknowledgments

None

## References

1. Provencio I, Rodriguez IR, Jiang G, Hayes WP, Moreira EF, Rollag MD. A novel human opsin in the inner retina. J Neurosci. 2000;20:600–605.

2. Berson DM, Dunn FA, Takao M. Phototransduction by retinal ganglion cells that set the circadian clock. Science. 2002;295:1070–1073.

3. Aranda ML, Schmidt TM. Diversity of intrinsically photosensitive retinal ganglion cells: circuits and functions. Cell Mol Life Sci. 2021;78:889–907.

4. Johnson J, Wu V, Donovan M, et al. Melanopsin-dependent light avoidance in neonatal mice. Proceedings of the National Academy of Sciences of the United States of America. 2010;107:17374–17378.

5. Delwig A, Logan AM, Copenhagen DR, Ahn AH. Light evokes melanopsin-dependent vocalization and neural activation associated with aversive experience in neonatal mice. PLoS One. 2012;7:e43787.

6. Bourin M, Hascoet M. The mouse light/dark box test. European Journal of Pharmacology. 2003;463:55–65.

7. Semo M, Gias C, Ahmado A, et al. Dissecting a role for melanopsin in behavioural light aversion reveals a response independent of conventional photoreception. PLoS One. 2010;5:e15009.

8. Matynia A, Parikh S, Chen B, et al. Intrinsically photosensitive retinal ganglion cells are the primary but not exclusive circuit for light aversion. Experimental eye research. 2012;105:60–69.

9. Matynia A, Parikh S, Deot N, et al. Light aversion and corneal mechanical sensitivity are altered by intrinscally photosensitive retinal ganglion cells in a mouse model of corneal surface damage. Experimental eye research. 2015;137:57–62.

10. Tamayo E, Mouland JW, Lucas RJ, Brown TM. Regulation of mouse exploratory behaviour by irradiance and cone-opponent signals. BMC Biol. 2023;21:178.

11. Okamoto K, Thompson R, Tashiro A, Chang Z, Bereiter DA. Bright light produces Fos-positive neurons in caudal trigeminal brainstem. Neuroscience. 2009;160:858–864.

12. Noseda R, Kainz V, Jakubowski M, et al. A neural mechanism for exacerbation of headache by light. Nature neuroscience. 2010;13:239–245.

13. Sowers LP, Wang M, Rea BJ, et al. Stimulation of Posterior Thalamic Nuclei Induces Photophobic Behavior in Mice. Headache. 2020;60:1961–1981.

14. Drummond PD. Photophobia and autonomic responses to facial pain in migraine. Brain : a journal of neurology. 1997;120 (Pt 10):1857–1864.

15. Boulloche N, Denuelle M, Payoux P, Fabre N, Trotter Y, Geraud G. Photophobia in migraine: an interictal PET study of cortical hyperexcitability and its modulation by pain. Journal of neurology, neurosurgery, and psychiatry. 2010;81:978–984.

16. Kowacs PA, Piovesan EJ, Werneck LC, et al. Influence of intense light stimulation on trigeminal and cervical pain perception thresholds. Cephalalgia. 2001;21:184–188.

17. McAdams H, Kaiser EA, Igdalova A, et al. Selective amplification of ipRGC signals accounts for interictal photophobia in migraine. Proceedings of the National Academy of Sciences of the United States of America. 2020;117:17320–17329.

18. Kaiser EA, McAdams H, Igdalova A, et al. Reflexive Eye Closure in Response to Cone and Melanopsin Stimulation: A Study of Implicit Measures of Light Sensitivity in Migraine. Neurology. 2021;97:e1672–e1680.

19. Selby G, Lance JW. Observations on 500 cases of migraine and allied vascular headache. Journal of neurology, neurosurgery, and psychiatry. 1960;23:23–32.

20. Drummond PD. A quantitative assessment of photophobia in migraine and tension headache. Headache. 1986;26:465–469.

21. Rasmussen BK, Jensen R, Olesen J. A population-based analysis of the diagnostic criteria of the International Headache Society. Cephalalgia. 1991;11:129–134.

22. Russell MB, Rasmussen BK, Fenger K, Olesen J. Migraine without aura and migraine with aura are distinct clinical entities: a study of four hundred and eighty-four male and female migraineurs from the general population. Cephalalgia. 1996;16:239–245.

23. Noseda R, Burstein R. Advances in understanding the mechanisms of migraine-type photophobia. Curr Opin Neurol. 2011;24:197–202.

24. Main A, Dowson A, Gross M. Photophobia and phonophobia in migraineurs between attacks. Headache. 1997;37:492–495.

25. Vanagaite J, Pareja JA, Storen O, White LR, Sand T, Stovner LJ. Light-induced discomfort and pain in migraine. Cephalalgia. 1997;17:733–741.

26. Vingen JV, Sand T, Stovner LJ. Sensitivity to various stimuli in primary headaches: a questionnaire study. Headache. 1999;39:552–558.

27. Shepherd AJ. Visual contrast processing in migraine. Cephalalgia. 2000;20:865–880.

28. Cady RK, Vause CV, Ho TW, Bigal ME, Durham PL. Elevated saliva calcitonin gene-related peptide levels during acute migraine predict therapeutic response to rizatriptan. Headache. 2009;49:1258–1266.

29. Goadsby PJ, Edvinsson L, Ekman R. Vasoactive peptide release in the extracerebral circulation of humans during migraine headache. Ann Neurol. 1990;28:183–187.

30. Gallai V, Sarchielli P, Floridi A, et al. Vasoactive peptide levels in the plasma of young migraine patients with and without aura assessed both interictally and ictally. Cephalalgia. 1995;15:384–390.

31. Bellamy JL, Cady RK, Durham PL. Salivary levels of CGRP and VIP in rhinosinusitis and migraine patients. Headache. 2006;46:24–33.

32. Recober A, Kuburas A, Zhang Z, Wemmie JA, Anderson MG, Russo AF. Role of calcitonin gene-related peptide in light-aversive behavior: implications for migraine. J Neurosci. 2009;29:8798–8804.

33. Recober A, Kaiser EA, Kuburas A, Russo AF. Induction of multiple photophobic behaviors in a transgenic mouse sensitized to CGRP. Neuropharmacology. 2010;58:156–165.

34. Kaiser EA, Kuburas A, Recober A, Russo AF. Modulation of CGRP-induced light aversion in wild-type mice by a 5-HT(1B/D) agonist. J Neurosci. 2012;32:15439–15449.

35. Mason BN, Kaiser EA, Kuburas A, et al. Induction of Migraine-Like Photophobic Behavior in Mice by Both Peripheral and Central CGRP Mechanisms. J Neurosci. 2017;37:204–216.

36. Estevez O, Spekreijse H. The “silent substitution” method in visual research. Vision research. 1982;22:681–691.

37. Mangutov E, Dripps I, Siegersma K, et al. Activation of delta-opioid receptors blocks allodynia in a model of headache induced by PACAP. British journal of pharmacology. 2025.

38. Patterson SS, Neitz M, Neitz J. S-cone circuits in the primate retina for non-image-forming vision. Semin Cell Dev Biol. 2022;126:66–70.

39. Guler AD, Ecker JL, Lall GS, et al. Melanopsin cells are the principal conduits for rod-cone input to non-image-forming vision. Nature. 2008;453:102–105.

40. Spitschan M, Jain S, Brainard DH, Aguirre GK. Opponent melanopsin and S-cone signals in the human pupillary light response. Proceedings of the National Academy of Sciences of the United States of America. 2014;111:15568–15572.

41. Matynia A, Nguyen E, Sun X, et al. Peripheral Sensory Neurons Expressing Melanopsin Respond to Light. Front Neural Circuits. 2016;10:60.

42. Gaskill BN, Gordon CJ, Pajor EA, Lucas JR, Davis JK, Garner JP. Heat or insulation: behavioral titration of mouse preference for warmth or access to a nest. PLoS One. 2012;7:e32799.

